# General Transcription Factor from *E. coli* with a Distinct Mechanism of Action

**DOI:** 10.1101/2023.09.17.558134

**Authors:** Nikita Vasilyev, Mengjie MJ Liu, Vitaly Epshtein, Ilya Shamovsky, Evgeny Nudler

## Abstract

Gene expression in *E. coli* is controlled by well-established mechanisms that activate or repress transcription. Here, we identify CedA as an unconventional transcription factor specifically associated with the RNA polymerase (RNAP) σ^70^ holoenzyme. Structural and biochemical analysis of CedA bound to RNAP reveal that it bridges distant domains of β and σ^70^ subunits to stabilize an open-promoter complex. Remarkably, CedA does so without contacting DNA. We further show that *cedA* is strongly induced in response to amino acid starvation, oxidative stress, and aminoglycosides. CedA provides a basal level of tolerance to these clinically relevant antibiotics, as well as to rifampicin and peroxide. Finally, we show that CedA modulates transcription of hundreds of bacterial genes, which explains its pleotropic effect on cell physiology and pathogenesis.

**One sentence summary:** An integrated structure-function approach uncovers CedA as a general transcription initiation factor in *E. coli* and elucidates its multifaceted role and unique mechanism.

## Introduction

*E. coli* RNA polymerase (RNAP) core enzyme consisting of five subunits (two identical α, β, β′ and ⍵) is capable of elongating growing RNA chain ^1^. At different transcription steps, RNAP core forms transient complexes with various accessory proteins. The initiation step is directed by σ factors responsible for sequence-specific recognition of promoter DNA and strands separation ^2,3^. Depending on environmental conditions, transcription initiation is finely tuned by a myriad of sequence-specific transcription factors ^4^ and nucleoid-associated proteins ^5^.

Besides its primary function in gene expression, RNAP is intimately involved in other essential cellular processes, such as DNA replication and repair ^6,7^. Negative supercoiling of DNA, resulting from active transcription in the vicinity of the origin of replication, promotes the binding of a replication initiation protein DnaA. Furthermore, a potential interaction between *E. coli* CedA and RNAP ^8–10^ ties transcription to another essential process, cell division ^11^. However, the nature of this connection remains poorly understood.

Gene encoding CedA was originally discovered as a multi-copy suppressor of the dnaAcos phenotype ^11^. At non-permissible temperature, mutant DnaA causes an over-initiation of DNA replication leading to non-dividing filamentous bacteria. Expression of *cedA* from a multi-copy plasmid restores *E. coli* capacity to divide. Based on this observation, CedA was ascribed as an activator of cell division.

*cedA* was found among the genes positively selected in uropathogenic *E. coli* strains ^12^. The ability of *E. coli* to form non-dividing filaments associated with persistent urinary tract infections ^13,14^ makes a connection to the original discovery of *cedA* as a suppressor of filamentous phenotype, implying that CedA plays a role in *E. coli* pathogenesis. Another clinically relevant observation was made in the study of bacterial response to gold nanoparticle-based antibiotics ^15,16^: CedA overexpression enhances *E. coli* tolerance to a compound named LAL-32, whereas the inactivation of a chromosomal copy of the gene results in higher sensitivity to this compound ^16^.

Giving its importance for cell division and clinical relevance, several groups set their efforts to identify the biological role and mechanism of CedA. Owing to its small size, CedA was a good target for the structural analysis by NMR ^9,17^. Structure of its C-terminal domain appeared similar to known structures of double-stranded DNA binding domains, whereas its N-terminus was largely unstructured ^17^. Indeed, DNA binding site of CedA was identified as TTTTXXT[T/G] using SELEX ^9^. However, the biological function and mechanism of CedA remain unknown, which prompted us to address these questions using a combination of quantitative proteomics, Cryo-EM, NGS, and biochemical approaches.

## Results

### CedA binds RNAP holoenzyme *in vivo*

Our approach to study protein-protein interactions *in vivo* uses immunoprecipitation (IP) of β′ subunit combined with mass spectrometry (MS)-based analysis of co-isolated proteins. We used *E. coli* strains bearing chromosomally 3×FLAG-tagged β′ (RpoC) and CedA to ensure natural levels of protein expression, and used crosslinking with formaldehyde to preserve *in vivo* composition of protein complexes ^18^.

Proteomic analysis revealed that nearly one thousand proteins consistently co-isolate with RNAP from exponentially growing cells (Fig. 1a). To identify true interactors, we ranked proteins by their abundance, however, even at a relatively high threshold cut-off (2% top-most abundant), less than half of the proteins were RNAP core subunits or known interactors (Fig. 1b and Supplementary Table 1). Many of the highly abundant proteins were ribosomal proteins and translation factors, which we reasoned could bind to the beads nonspecifically and rank highly due to their sheer abundance in bacteria. At the same time, RNAP-interacting proteins expressed at low levels would automatically rank low. CedA, which abundance was measured at 235 copies per cell ^19^ (approximately 10% that of RpoC) is placed at position 129 in the list of top-most abundant proteins co-isolated with RpoC (Supplementary Table S1), and would not be considered RNAP-interacting protein. To mitigate these issues, we introduced two additional metrics. First, we performed the pulldown experiments for bacteria bearing no FLAG-tagged proteins to calculate a specificity rank proportional to the difference in abundance between target and non-specific pull downs. Secondly, we measured the level of proteins in a lysate and calculated the enrichment rank proportional to the difference in the abundance between lysates and a target pulldown. With all the metrics combined, 19 of 20 proteins among the top-most ranking 2% were known RNAP interactors (Fig. 1d and Supplementary Table 2), showing a significant improvement in sorting out non-specific candidates. Now, CedA is among top-most ranking proteins at position 10, following RNAP core subunits, sigma factors RpoD and FecI, and elongation/termination/antitermination factors NusA, NusG and SuhB (Supplementary Table 2).

**Fig. 1.**
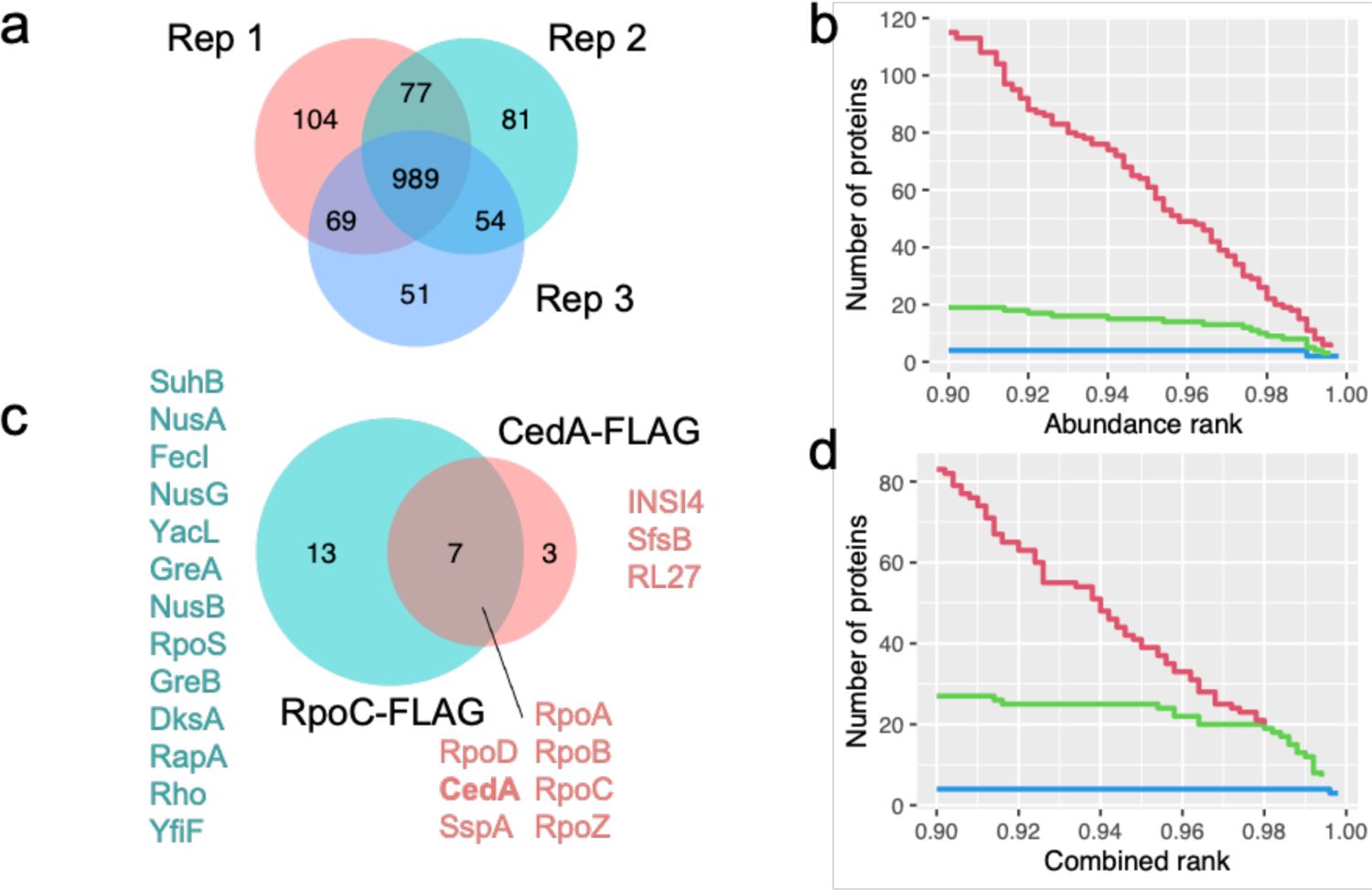
RNAP and CedA interactomics in *E. coli*. **a**, Venn diagram showing an overlap of proteins co-isolated with RNAP (FLAG-tagged RpoC) in three independent experiments. **b**, Step plot showing a number of proteins identified at a given abundance rank threshold. Plotted are top 10% most-abundant (rank 0.9 – 1.0) proteins. Colors represent RNAP core subunits (blue), known RNAP interactors (green), other *E. coli* proteins (red). **c**, Venn diagram showing distinct and common proteins co-isolated with both RNAP (FLAG-tagged RpoC) and CedA (FLAG-tagged) identified at a 2% combined rank threshold. **d**, Step plot showing a number of proteins identified at a given combined rank threshold (top 10% proteins plotted). Colors are the same as in (B).

Using this improved procedure, we performed the same measurements for CedA, and when the top-most ranking 2% of the proteins co-isolated with CedA were compared to those isolated with RNAP, seven were common between both targets (Fig. 1c, Supplementary Tables 2 and 3). These proteins were RNAP core subunits (RpoA, RpoB, RpoC, and RpoZ), CedA, transcription initiation factor σ^70^ (RpoD), and stringent starvation protein A (SspA). This result implied a direct involvement of CedA with σ^70^-mediated transcription initiation.

### CedA resides at σ^70^ promoters

CedA was reported to bind DNA, presumably via its C-terminal domain, which is structurally similar to known double-stranded DNA binding proteins ^17^. The SELEX study ^9^, identified a consensus binding sequence for CedA as a short T-rich stretch of DNA. These reports, together with our initial observation of CedA affinity to RNAPσ^70^ holoenzyme, led us to hypothesize that CedA could be a sequence-specific transcription initiation factor. To test this hypothesis, we performed a ChIP-seq experiment to identify CedA binding sites on *E. coli* chromosome.

We isolated RNAP and CedA from formaldehyde-treated bacteria by IP and analyzed the coprecipitated DNA using the high-throughput sequencing. Sequencing depth profiles demonstrate that CedA locates preferentially at promoter regions, whereas the core RNAP signal is distributed along entire transcription units (Fig. 2a). To identify a consensus sequence of a CedA-binding site, we took 20-bp long DNA sequences centered around CedA peaks and aligned them using MAFFT^20^. The resulting sequence (Fig. 2b) matches the σ^70^ promoter −10 consensus TATAAT ^21^. Further analyzes revealed that the majority of identified CedA peaks (301 of 473) are indeed located around −10 region of *E. coli* promoters, and map within the 20-bp distance from known transcription start sites (Fig. 2c). Among the 172 unassigned CedA peaks, 117 were mapped to the intergenic regions, implying their possible association with yet uncharacterized promoters and transcription start sites (Supplementary Table 4).

**Fig. 2.**
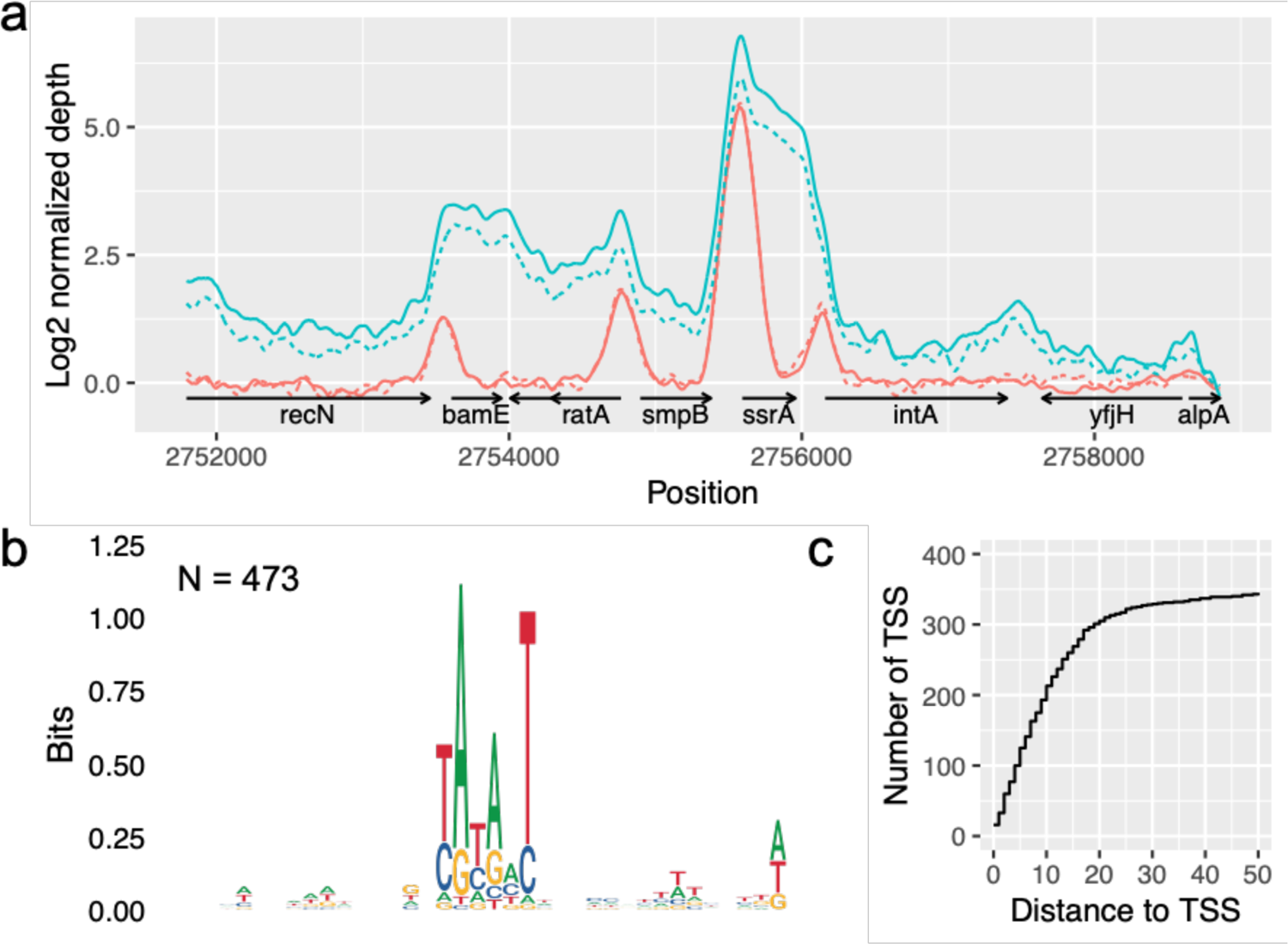
CedA colocalizes with the RNAPσ^70^ holoenzyme throughout *E. coli* genome. **a**, Smoothed ChIP-seq sequence depth profile of RNAP (FLAG-tagged RpoC, cyan) and CedA (FLAG-tagged CedA, red). Solid and dashed lines represent two replicates of the experiment. **b**, Consensus DNA sequence identified by aligning 20-bp long DNA segments centered around CedA peaks. **c**, Step plot showing a number of transcription start sites (TSS) mapped at various distances from CedA peaks.

A combination of high-throughput proteomics and NGS provided strong evidence supporting the involvement of CedA in transcription initiation: CedA binds to RNAPσ^70^ holoenzyme and is localized around the −10-consensus sequence of σ^70^ promoters. It associates with promoters of hundreds of genes involved in every essential process, including transcription, translation, DNA replication, repair, recombination, transport and metabolism of nucleotides, amino acids, lipids, carbohydrates, and cofactors; cell wall synthesis, cell division, and others (Supplementary Table 4).

### Structure of CedA bound to the open promoter complex

To gain further insight into the mechanism of CedA, we employed cryo-EM to solve the structure of CedA bound to RNAPσ^70^-open promoter complex. Purified RNAPσ^70^ holoenzyme was first incubated with 85-bp DNA representing −60 to +25 positions relative to the transcription start site (TSS) of the *ssrA* promoter (Fig. 3a), which has the strongest association with CedA, as determined by ChIP-seq (Fig. 2a and Supplementary Table 4). Purified CedA was added to the pre-formed RNAPσ^70^•ssrAp and the resulting complex was re-purified by gel-filtration and used for the single particle cryo-EM analysis. A structure of the whole complex was determined at the overall resolution of 2.76 Å (Extended data Fig. 1), which let us to build CedA model *de novo*.

**Fig. 3.**
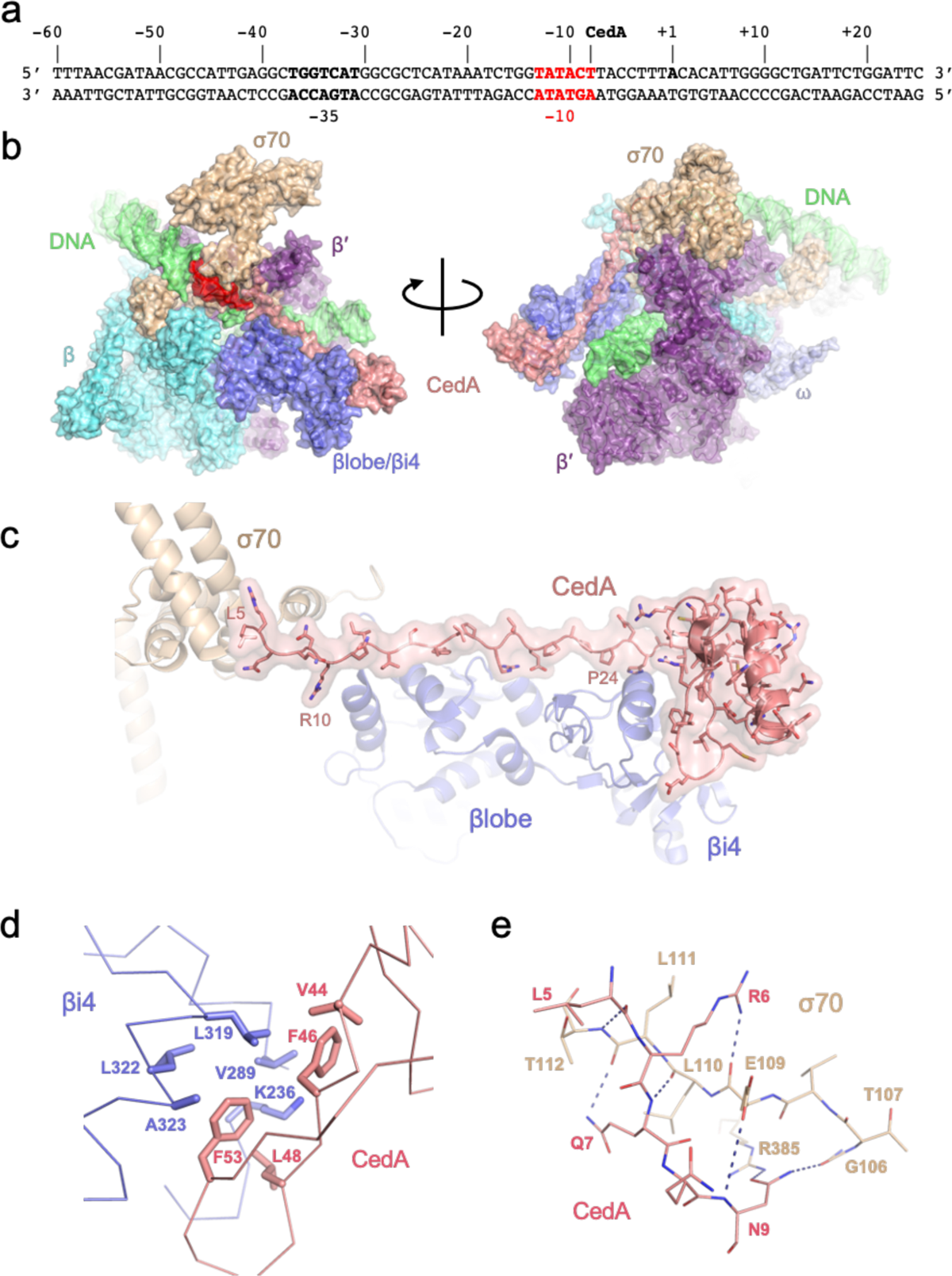
Cryo-EM structure of the CedA-bound open promoter complex. **a**, DNA sequence of ssrAp promoter. Position of CedA peak is marked, and −10 consensus sequence is shown in red. **b**, Surface representation of CedA-bound OPC shown from two angles. Colors represent individual subunits and their parts: DNA (green), β subunit (cyan), βlobe/βi4 (dark blue), β′ subunit (purple), ⍵ subunit (light blue), σ70 (wheat), CedA (salmon). **c**, Close-up view of CedA interaction with βlobe/βi4 and σ70. **d**, Interface between CedA C-terminal domain and βi4 formed by hydrophobic residues. **e**, Hydrogen bonds formed between N-terminus of CedA and σ70 region 1.2.

A general architecture of RNAP in the complex (Fig. 3b) is similar to the known structure of a transcription initiation complex ^22^. We identified CedA adopting a ladle-like shape and binding the RNAP at the tip of a β-pincer formed by a lineage-specific sequence insertion βi4, β-lobe, and σ^70^ (Fig. 3b-e).

Unexpectedly, the C-terminal globular domain of CedA (residues 27–80), structurally similar to dsDNA-binding proteins ^17^, does not contact DNA. Instead, it binds βi4 via predominantly hydrophobic contacts (Fig. 3d). An N-terminal part of CedA, which was reported as unstructured ^9,17^, stretches along the DNA-facing side of β-lobe, and the very N-terminus (residues 5–10), bridging the gap between the β-lobe and σ^70^ region 1.2 (Fig. 3c, 3e), and effectively locking the melted DNA in the RNAP main channel.

The N-terminus of CedA forms several hydrogen bonds with σ^70^ (Fig. 3d). The only side-chain-specific contact was found between CedA Asn9 and σ^70^ Arg385, whereas other contacts involve main-chain atoms (Fig. 3e). A sequence alignment of *E. coli* σ factors (Extended data Fig. 2) reveals a high degree of similarity between σ^70^ and σ^S^ in region 1.2, while σ^32^ appears more distant, and other σ factors have no apparent similarity. A structural alignment (Extended data Fig. 3) mirrors the sequence comparison, implying that CedA could recognize RNAP holoenzyme bound to σ^S^, although, strength of the interaction would be diminished due to the loss of a contact between CedA Asn9 and σ^70^ Glu109 replaced by a proline in σ^S^ and σ^32^ (Extended data Fig. 3).

A close spatial proximity of the CedA N-terminus to σ^70^ regions 1.1 and 2, which are bound to a non-template DNA strand, explains why an apparent DNA binding site of CedA was identified as a −10 σ^70^ promoter consensus sequence. In spite of the lack of any direct contacts between CedA and DNA, formaldehyde crosslinking would tie CedA to the σ^70^-bound −10 consensus DNA.

A closer comparison of our structure with other models of the IC revealed a major difference with DksA/TraR-bound structures ^23,24^. DksA, and its homolog TraR, bind in the secondary channel of RNAP affecting kinetics of transcription initiation in a promoter sequence-dependent manner ^25,26^. Upon binding to the IC, these proteins cause a major conformational change involving the movement of β-lobe/βi4 to form contacts with DksA (or TraR) (Fig. 4a), which widens the main channel of RNAP, thus facilitating a reversal of the open-promoter complex (RPo) ^23,24^. This conformational change appears to be incompatible with our structure, as CedA would prevent such movement by keeping β-lobe/βi4 tied to σ^70^ (Fig. 3c). This observation predicts that CedA has an overall stabilizing effect on RPo by keeping DNA melted in the major groove of RNAP and, thus, also counteracting the effect of DksA (or TraR).

**Fig. 4.**
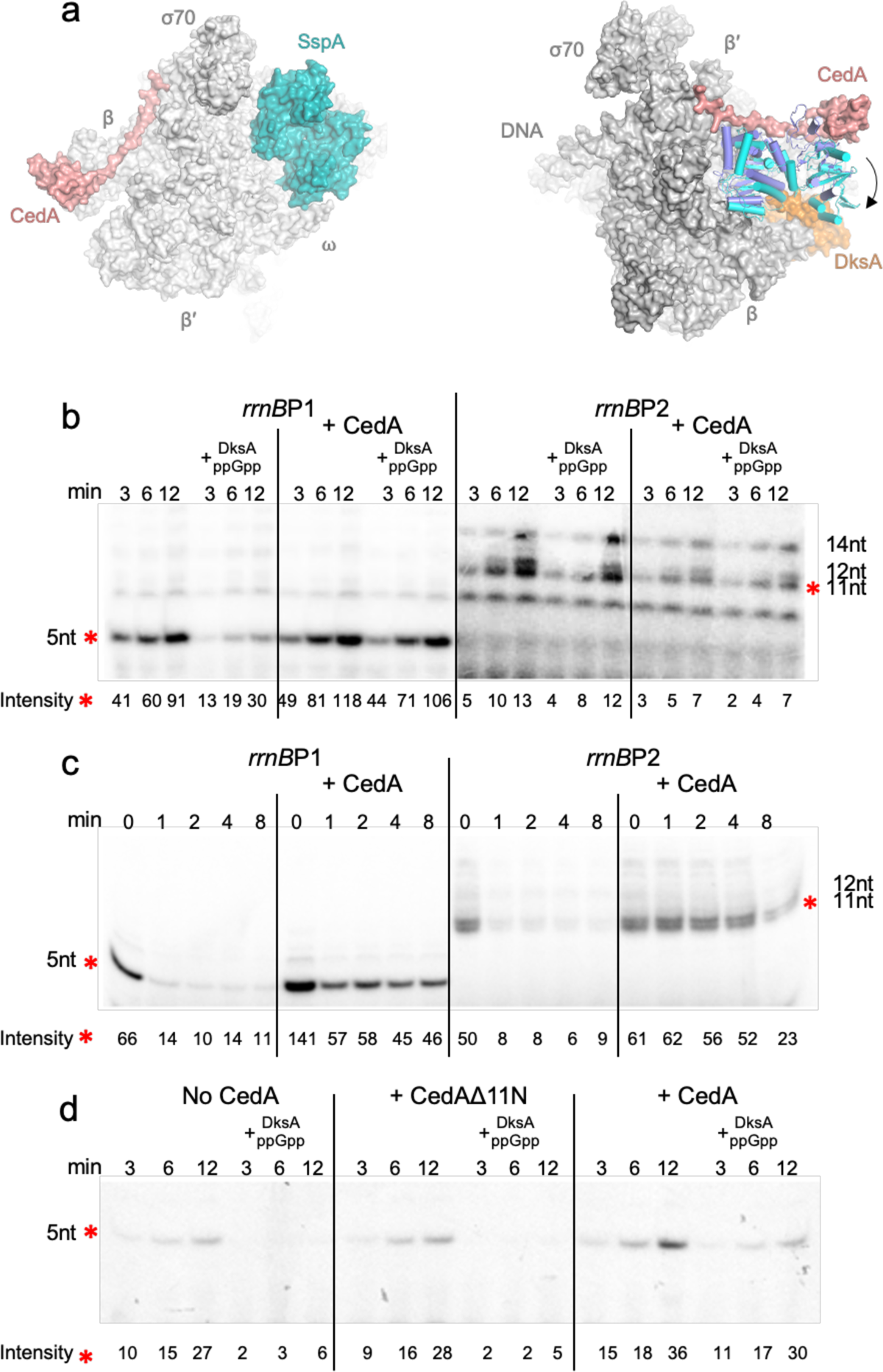
Modulation of transcription initiation by CedA. **a**, Structural comparison of CedA-, SspA- and DksA-bound open promoter complex. Structures of SspA-(PDB: 7C97) ^48^ (left) and DksA-bound (PDB: 7KHI) (right) RPo were aligned with CedA-bound structure using PyMol 2.4. SspA and DksA are shown in teal and orange, respectively. Conformational change in β-lobe/βi4 upon DksA binding is shown by an arrow. Structure of β-lobe/βi4 in the DksA-bound complex is shown in cyan, CedA-bound complex–dark blue. **b-d**, Representative autoradiograms of the RNA products from the transcription initiation assay utilizing *rrnB*P1 or *rrnB*P2 promoters. Relative intensity values for the bands marked with red asterisks were obtained using ImageQuant software (see Methods). Normalization between independent experiments was achieved by using the bands with highest intensity readings. SD was calculated from at least three independent experiments. **b**, Reactions were incubated for 3, 6, and 12 min in the presence of DksA/ppGpp and/or CedA. **c**, After the assembly of the open-promoter complex, reactions were either immediately supplemented with NTPs (time point 0) or mixed with heparin and incubated for 1, 2, 4 and 8 min before the addition of NTPs. **d**, Reactions initiated from *rrnB*P1 were incubated for 3, 6 and 12 min with or without DksA/ppGpp together with CedA or its truncated version lacking the eleven N-terminal amino acids (CedAΔ11N).

### CedA stabilizes an open-promoter complex and counteracts DksA

To test our structure-based predictions, we performed *in vitro* transcription reactions using DNA templates containing DksA-sensitive and -resistant promoters of the *E. coli* ribosomal operon *rrnB*. To investigate a potential effect of CedA on transcription initiation, the reactions were performed with a limited set of nucleotides (ATP and CTP only) allowing the synthesis of short RNA products before the transition to a productive elongation phase.

The transcriptional assay in the presence of DksA and its co-effector ppGpp confirmed the inhibition of DksA-sensitive promoter *rrnB*P1, and no effect on *rrnB*P2 (Fig. 4b). When CedA was present, DksA/ppGpp were no longer able to inhibit *rrnB*P1 transcription, whereas *rrnB*P2 transcription was mildly suppressed by CedA independently of DksA/ppGpp. These results validate our prediction of CedA counteracting DksA, and also showed that CedA may have its own IC inhibitory activity. This inhibitory effect of CedA may result from an increased lifetime of RPo, leading to a delayed promoter clearance and, as a result, a reduced reaction turnover rate. Next, we compared the effect of CedA on the RPo lifetime. Following the RPo assembly, transcription reactions were incubated with heparin, a non-specific competitor of DNA that prevents re-initiation of transcription. After various times of incubation with heparin, the reactions were chased with NTPs to allow for a single-round RNA synthesis (Fig. 4c). CedA affected both promoters in a positive way resulting in the increased amount of synthesized RNA after the incubation with heparin. This result shows that CedA does indeed increase the lifetime of RNAP-promoter complex by preventing it from dissociation, hence the increased resistance to heparin. Finally, to verify that the observed effects were due to constrains imposed by CedA on RNAP plasticity at the promoter, we examined a CedA mutant (CedAΔ11N) lacking the first eleven amino acids. Based on the structure, this truncated mutant should not bind σ^70^ and, hence, cannot prevent β-lobe/βi4 from the DksA-induced conformational change. As predicted, CedAΔ11N failed to counteract DksA-mediated inhibition of transcription at *rrnB*P1 (Fig. 4d).

### CedA renders *E. coli* more tolerant to oxidative stress and selected antibiotics

CedA is encoded by a single-gene operon. Its promoter is directly adjacent to *katE* promoter, facing the opposite direction (Fig. 5a). KatE (HPII) is one of the two catalases in *E. coli*, which expression depends on general stress sigma factor α^S^ ^27^. The direct proximity between *katE* and *cedA* promoters suggests the two genes may share a common function. Indeed, the expression of *katE* and *cedA* is induced approximately 6-fold by peroxide (Fig. 5b, b). Furthermore, cells overexpressing CedA became more resistant to peroxide than control cells (Fig. 5d).

**Fig. 5.**
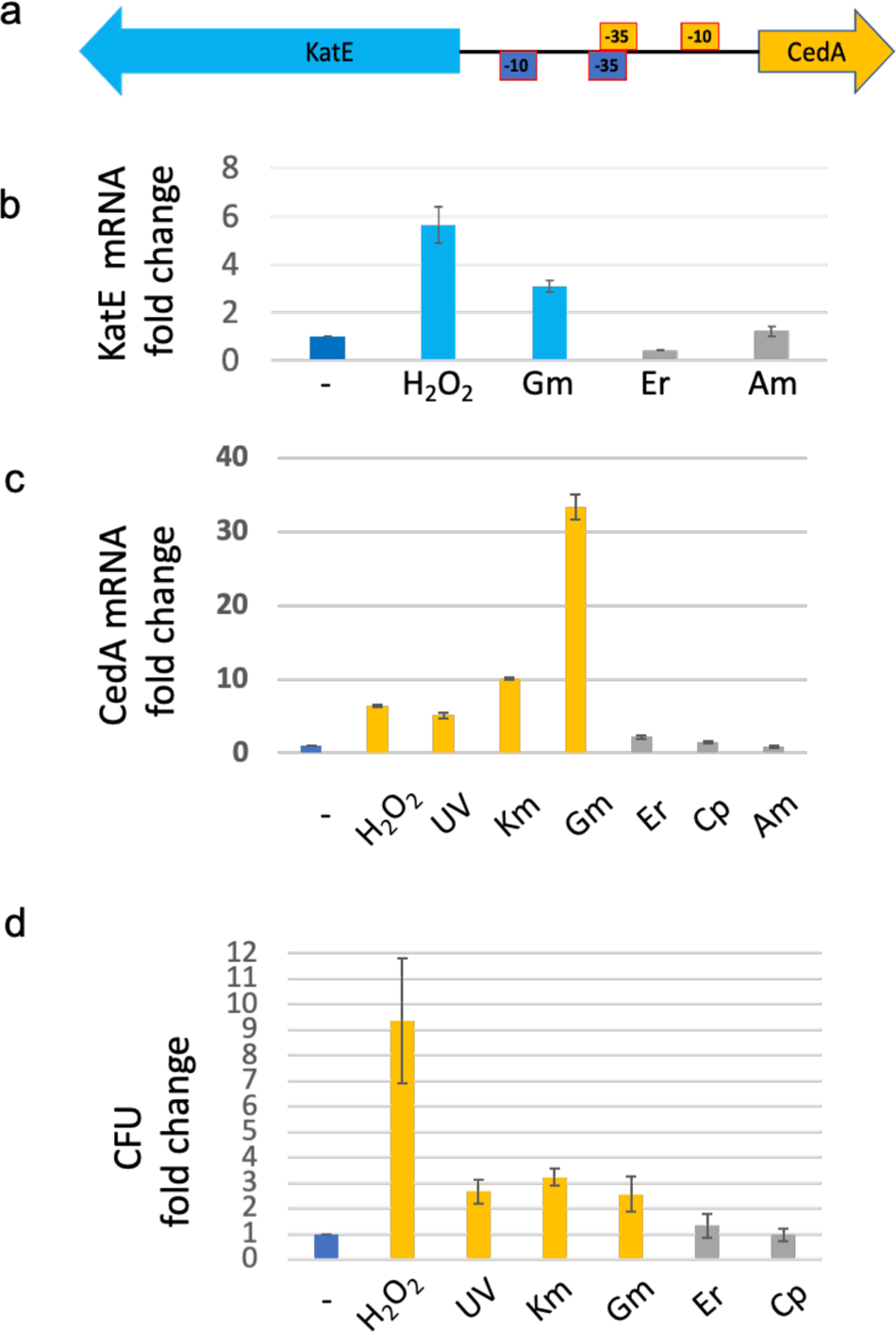
CedA renders *E. coli* cells more tolerant to peroxide and aminoglycosides. **a**, Schematics of *cedA* promoter region. Arrows indicate the direction of transcription. The −10 and −35 elements of *katE* and *cedA* promoters are colored in blue and yellow, respectively. **b** and **c**, *katE* and *cedA* mRNA changes in response to different stress conditions as determined by RT-qPCR. The amounts of *katE* and *cedA* mRNA were normalized to that of *gapA*. The resulting changes in the reaction threshold cycle values (ΔCt) were compared to non-stressed control (dark blue bars). −log2 differences in Ct values (ΔΔCt) were plotted as fold changes. Significant changes (P < 0.01) are shown as blue (**b**) or yellow (**c**) bars, not significant changes—gray bars. H_2_O_2_ – 2 mM, UV – 64 J cm^-2^, Km – kanamycin (50 µg/ml), Gm – gentamycin (20 µg/ml), Er – erythromycin (50 µg/ml), Cp – ciprofloxacin (1 µg/ml), Am – ampicillin (100 µg/ml). **d**, Survival rate of *E. coli* cells treated with the stressors as in (c) following the plasmid-born *cedA* induction. Significant changes in CFUs (P < 0.05) are shown as yellow bars, not significant changes—gray bars. Values (fold change) were normalized to an empty vector control and compared to a non-stressed condition (dark blue bar). The experiments (b-d) were repeated at least trice. RT-qPCR was performed using three independent biological replicates with three technical replicates for each pair of primers.

As many antibiotics promote oxidative stress ^28^, we also examined whether CedA offers any protection against different classes of antibiotics. We found that cells overexpressing CedA become more resistant to aminoglycosides gentamycin and kanamycin (Fig. 5d). CedA appears to have no effect on cellular tolerance to quinolones (ciprofloxacin) and macrolides (erythromycin) (Fig. 5d). Concordantly, aminoglycosides, but not erythromycin or ciprofloxacin, also stimulated *cedA* transcription (Fig. 5c).

The colony forming ability of Δ*cedA* cells was greatly diminished in the presence of RNAP inhibitor rifampicin (Fig. 6a). A plasmid overexpressing full-length CedA, but not its truncated version, not only abolished rifampicin sensitivity of Δ*cedA* cells, but also let them to grow at higher concentrations of rifampicin comparing to the wt, highlighting CedA involvement in rifampicin tolerance as well.

**Fig. 6.**
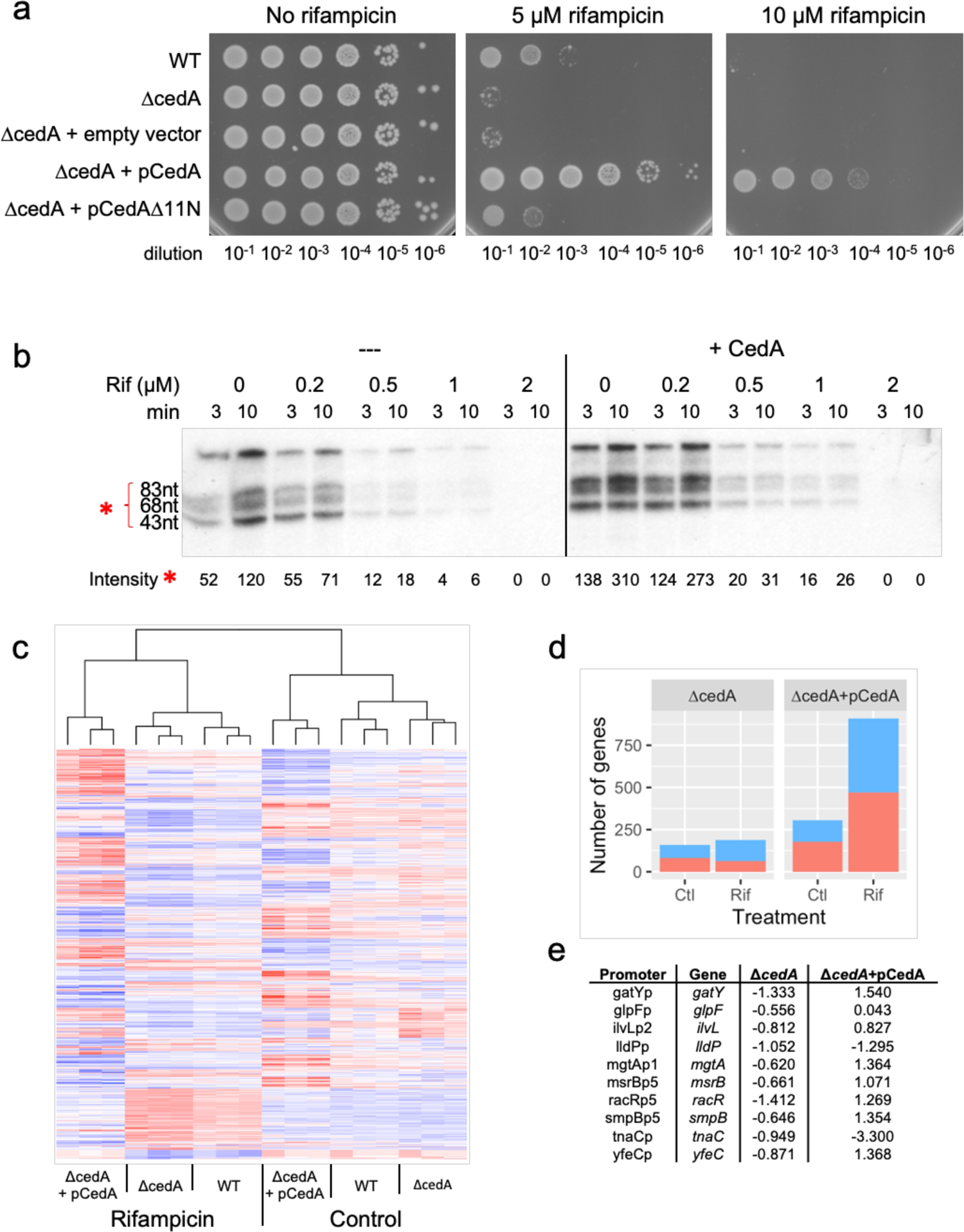
CedA promotes rifampicin tolerance and alters global gene expression in *E. coli*. **a**, Representative efficiencies of colony formation of parent wt, Δ*cedA*, or Δ*cedA* mutant transformed with plasmids bearing no insert (empty vector), full-length *cedA* (pCedA), or truncated *cedA* lacking the 5′-terminal 11 codons (pCedAΔ11N). Bacteria were grown overnight on LB agar plates containing 10 ng/ml anhydrotetracycline (inducer) and 0, 5, 10 μM rifampicin. **b,** Autoradiograms of the RNA products of multi-round transcription reactions performed in the presence of rifampicin and CedA using DNA template containing T7A1 promoter. Relative intensity values and SD for the bands marked with an asterisk were calculated as in Fig. 4 from three independent experiments. **c-e**, Differentially expressed *E. coli* genes regulated by CedA. **c**, Heatmap representation of a change in mRNA levels of differentially expressed genes (factorial ANOVA q-value < 0.01) in response to the *cedA* deletion or overexpression and treatment with rifampicin. Upregulated genes are shown in red, downregulated—in blue. **d**, Number of up-(red) and downregulated (blue) genes filtered by q-value below 0.01 and log2 fold change above 0.5. Log2 fold change was calculated for the deletion (Δ*cedA*) and overexpression (Δ*cedA*+pCedA) strains relative to the wt. **e**, List of promoters associated with CedA that responded to rifampicin treatment (q-value < 0.01, log2 fold change > 0.5) in Δ*cedA* strain. Columns Δ*cedA* and Δ*cedA*+pCedA show magnitude of change of gene expression levels (log2 fold change) relative to the wt in the presence of rifampicin in *cedA* deletion and overexpression strains, respectively.

To identify whether rifampicin tolerance resulted from a direct modulation of RNAP activity by CedA, we performed transcription reactions in the presence of rifampicin using a DNA template containing a generic σ^70^ promoter T7A1. CedA activated transcription resulting in a higher overall yield of RNA, however, degree of RNAP inhibition by rifampicin was not affected (Fig. 6b). This result suggests that CedA does not alter the mode of interaction between RNAP and rifampicin, but contributes to antibiotic tolerance by promoting residual transcription in the presence of the drug.

### CedA modulates hundreds of *E. coli* genes

As antibiotic tolerance may involve multiple differentially expressed genes responding to CedA, we performed RNA-seq experiment to compare gene expression between wt bacteria and strains lacking or overexpressing CedA.

The wt, Δ*cedA*, and Δ*cedA* cells transformed with pCedA were grown in LB at 37 °C to OD_600_∼0.15–0.2, before cultures were supplemented with rifampicin (10 μM). After 1-hour incubation, bacteria were harvested and used for RNA isolation. Differentially expressed genes responding to both CedA and rifampicin were identified by factorial ANOVA test using log2-transformed normalized reads counts as a measure of mRNA levels.

We identified 1,309 genes (q-value < 0.01) responding to CedA and rifampicin (Supplementary Table 5). The overall pattern of gene expression (Fig. 6c) shows that in both control and rifampicin conditions, the overexpression of CedA has a much larger overall effect, whereas the pattern of gene expression in Δ*cedA* mostly follows that of the wt, with many genes yet significantly affected. To further clarify the effect of CedA, we filtered differentially expressed genes based on the magnitude of the response, keeping only those with log2 fold change above 0.5 when compared to the wt. Both up- and downregulated genes represented a significant fraction in both strains (Fig. 6d), with a larger number of genes alternatively regulated in a strain overexpressing CedA.

Many genes affected by CedA were transcription factors, RNA modifying enzymes, and ribonucleases (Supplementary Table 5), suggesting that globally altered mRNA levels could be a result of an indirect action of CedA. Thus, to identify direct effects of the protein, our next step was to look at the effect of CedA on genes which promoters were associated with it based on ChIP-seq (Supplementary Table 4). All of the 10 promoters that passed the filtering criteria (q-value < 0.01, log2 fold change above 0.5) were downregulated in rifampicin-treated Δ*cedA* cells (Fig. 6e). Overexpression of CedA effectively reversed the inhibitory effect of rifampicin for eight of them, bringing the expression level of downstream genes to that of the wild type or above.

Results of RNA-seq experiment show that CedA modulates gene expression in condition-dependent and promoter-dependent manner. An apparent transcriptional repression in cells grown without the antibiotic may result from the increased lifetime of RPo leading to delayed promoter clearance, as we proposed to explain the inhibitory effect of CedA on multi-round transcription directed by *rrnB*P2 (Fig. 4b). Stimulation of transcription by CedA appears to be crucial for rifampicin tolerance and potentially involves promoters that are particularly sensitive to the antibiotic.

## Discussion

In this study, we combined chemical crosslinking and affinity isolation followed by MS-based proteomics and next generation sequencing to identify RNAP-associated protein CedA as a global transcription initiation factor.

Our experimental setup relies on *in vivo* crosslinking with formaldehyde, a fast acting and robust crosslinking agent, to fix protein-protein complexes in their natural environment prior to affinity isolation and analysis by MS. A typical experiment would identify hundreds of proteins, among which only 5-10% top ranking may be considered as true interactors. Proteins that bind to affinity media non-specifically were excluded from the analysis as false positives. In addition to ranking proteins by their abundance, we measured degree of specificity and enrichment, which all together were summarized in a combined rank, greatly improving the identification of RNAP interacting partners. With the same analysis applied to CedA, we were able to confidently call RNAPσ^70^ holoenzyme as the primary *in vivo* interacting partner of CedA. A complementary ChIP-seq analysis identified σ^70^ promoters as a target for CedA in *E. coli* genome. Overall, a combination of high-throughput methods, as exemplified in this study, provides an efficient way to determine interacting macromolecules and infer their function.

To understand the mechanism of CedA we used cryo-EM to solve the structure of CedA bound to the RPo. With the high-resolution map, we were able to trace most of the CedA, which acquires ladle-shaped conformation when bound to RNAP. To our surprise, CedA, which was predicted to bind DNA via its C-terminal domain ^9^, interacts exclusively with RNAP β subunit and σ^70^, and not with DNA. A cup of the ladle-shaped CedA molecule, the C-terminal globular domain, binds at the tip of β-pincer that is formed by βi4 of RNAP. Specificity of CedA for a lineage-specific sequence insertion explains why it is conserved in proteobacteria. The rest of the molecule, the elongated N-terminal part of CedA (handle of the ladle) stretches along β-lobe and binds to σ^70^ region 1.2. Sequence alignment of *E. coli* σ factors suggests that CedA may, in principle, also bind σ^S^ and σ^H^ holoenzymes. However, our proteomic analysis of CedA interactome does not support alternative σ as plausible interactors *in vivo*.

Based on our structural analysis, we predicted that CedA should have an overall stabilizing effect on RPo by restricting the movement of β-lobe, while keeping it in a closed conformation, contrasting an open conformation induced by DksA•ppGpp or TraR^23,24^. *In vitro* experiments confirmed our predictions. We found that CedA stimulates transcription from DksA-sensitive promoter *rrnB*P1 and has a stabilizing effect that increased the lifetime of RPo when challenged with heparin.

CedA renders *E. coli* more resistant to oxidative stress and antibiotics that inhibit translation (aminoglycosides) and transcription (rifampicin). Moreover, *cedA* is strongly induced in response to aminoglycosides and peroxide, demonstrating the involvement of CedA in bacterial tolerance. When tested *in vitro*, CedA activated residual transcription in the presence of rifampicin, but did not change RNAP sensitivity to the antibiotic.

RNA-seq analysis shows that CedA affects hundreds of genes under normal growth conditions. We identified over a thousand of genes that were affected by rifampicin treatment in CedA-dependent manner. The list is vast, which precludes us from pinpointing any specific candidate, or their combination, underlying rifampicin tolerance. However, we analyzed the expression of genes located immediately downstream of CedA-associated promoters identified by ChIP-seq to show that in cells overexpressing CedA these promoters were upregulated upon rifampicin treatment, whereas in the deletion strain they were suppressed. Thus, we conclude that the stimulation of transcription by CedA contributes to rifampicin tolerance. A similar global transcriptional response mediated by CedA is likely to underlie tolerance to other clinically relevant antibiotics we examined here and to oxidative stress, highlighting CedA as a new promising antimicrobial target, especially in light of its positive retention in uropathogenic *E. coli*^12^.

## Acknowledgments

We thank William Rice, Bing Wang, David Stokes and Zheng Liu for helping with sample screening and data collection at NYULH Cryo-EM Laboratory. We thank the BigPurple HPC core at NYULH for computer access. This work was supported by the NIH grant R01 GM126891, Blavatnik Family Foundation, and by the Howard Hughes Medical Institute (E.N.).

## Author Contributions

N.V. carried out all genetic, proteomics, and transcriptomics studies. M.L. purified the proteins, collected and processed EM data, and built the structural model. V.E. performed *in vitro* transcription assays. I.S. performed the NGS. E.N. supervised the project. N.V. and E.N. wrote the manuscript with input from all the authors.

## Materials and methods

### Bacterial strains and plasmids

*E. coli* MG1655 wt and its derivatives were used in ChIP-seq and RNA-seq experiments, analysis of protein interactions *in vivo*, and rifampicin resistance. *E. coli* MG1655 *rpoC*-3×FLAG was among the strains in laboratory collection ^29^, *E. coli* MG1655 *cedA*-3×FLAG and *E. coli* MG1655 Δ*cedA* were prepared in this study. *E. coli* BL21(DE3) and NEB Turbo Competent *E. coli* (NEB) were used for proteins overexpression and DNA cloning, respectively. paTc-based vectors ^29^ were used for tetracycline-inducible expression of CedA (paTc-*cedA*) and its truncated variant lacking first 11 amino acids (paTc-*cedA*Δ11N). Plasmids pVS10 ^30^ and pSumoH10-*cedA* were used for the overexpression of *E. coli* RNAP and CedA, respectively.

### Chromosomal deletions and tagging

The modification and/or deletion of chromosomal genes were done using λ Red recombinase system as described elsewhere^31^. To incorporate a 3×FLAG epitope at the C-terminus of a target protein, a kanamycin resistance cassette from pKD4 was first PCR-amplified using a pair of primers (Supplementary Table 6) introducing a 3×FLAG-coding region at the 5ʹ terminus of the cassette. Then a 3×FLAG-bearing cassette was amplified with a gene-specific pair of primers to introduce a 3×FLAG-encoding sequence at the 3ʹ end of a target gene immediately before the stop codon. For a gene deletion, a kanamycin resistance cassette was amplified with a pair of primers including the sequences flanking an open reading frame of a target gene. Resulting DNA fragments were introduced by electroporation into *E. coli* MG1655 bearing pKD46. After 2-hour recovery in SOC media at 30 °C, bacteria were plated on LB agar containing 50 μg/ml kanamycin and colonies were grown at 37 °C for 16-20 hours. Individual colonies were re-streaked on LB plates containing 50 μg/ml kanamycin, grown overnight at 37 °C and screened for loss of pKD46, which was confirmed by lack of ampicillin resistance when bacteria were grown on LB agar containing 100 μg/ml carbenicillin.

DNA from positive clones was extracted using a Monarch Genomic DNA Purification kit (NEB). DNA insertion was confirmed by amplification of a fragment including both genomic and kanamycin resistance cassette sequences followed by the sequencing of amplified DNA (Macrogen).

### Plasmid construction

Plasmids were constructed using a NEBuilder HiFi DNA Assembly kit (NEB) according to manufacturer instructions. Assembly reactions contained reverse-amplified vector DNA (paTc or pSumoH10) treated with Dpn I and a DNA fragment comprising of *cedA* gene amplified from *E. coli* MG1655 chromosome flanked by vector-specific sequences. After transformation, NEB Turbo *E. coli* were selected on LB agar containing appropriate antibiotic (100 μg/ml carbenicillin for paTc, 50 μg/ml kanamycin for pSumoH10). Individual clones were then grown in liquid LB, plasmid DNA was isolated using PureLink Plasmi Miniprep kit (Thermo) and correct inserts were confirmed by sequencing (Macrogen).

### In vivo crosslinking

*E. coli* MG1655 *rpoC*-3×FLAG or *cedA*-3×FLAG strains were grown overnight at 37 °C in LB media containing 50 μg/ml kanamycin. 0.3 ml of overnight cultures were inoculated into 30 ml of fresh LB contained in 125-ml flasks and new cultures were grown to mid-log phase (OD_600_ 0.4–0.6) at 37 °C with shaking at 250 rpm. For crosslinking, 16% methanol-free formaldehyde (Fisher) was added to 0.3% final concentration and the incubation continued for 10 min. To neutralize formaldehyde, 2 M glycine was added to cultures at 0.2 M final concentration for 5 min. Bacteria were then collected by 10-min centrifugation at 4,000g, 4°C. Pellets were resuspended in 0.3 ml of lysis buffer: 50 mM Tris-HCl pH 7.5, 150 mM NaCl, 10 mM MgCl_2_, 10% glycerol, 0.5% Triton X-100, and protease inhibitors cocktail (Roche), and stored at −20 °C. Lysis buffer for ChIP-seq experiments contained 1 mM EDTA instead of 10 mM MgCl_2_.

### Immunoprecipitation

Bacterial samples that were stored frozen as suspension in a lysis buffer were thawed on ice. Lysozyme (Roche) and Pierce Universal Nuclease (Thermo) were both added to 0.5 U/μl final concentration. To prevent DNA degradation in the ChIP samples, Pierce Universal Nuclease was replaced with 5 μg/ml RNase A. Bacteria were then disrupted by 20 cycles of sonication in Bioruptor Pico (Diagenode) with each cycle lasting for 30 sec between 30 sec breaks. In ChIP samples, the sonication ensured DNA fragmentation to an average size 150 bp. Lysates were clarified by 10-min centrifugation at 20000g, 4 °C and transferred to the new tubes. To capture FLAG-tagged proteins, Pierce Anti-DYKDDDDK Magnetic Agarose (Thermo) pre-equilibrated in the lysis buffer was added at 5 μl of packed beads per sample followed by 2-hour incubation at 4 °C on a rotary mixer. Beads were washed four times with a 1 ml ice-cold wash buffer (20 mM Tris-HCl pH 7.5, 500 mM NaCl, 1 mM EDTA, 0.1% Triton X-100) and kept on ice before proceeding to the next step of either on-beads digestion of proteins with trypsin for LC-MS analysis, or DNA extraction for the construction of sequencing libraries.

### On-beads proteins digestion

Following the immunoprecipitation procedure, beads were washed twice with 1 ml ice-cold NH_4_HCO_3_ to exchange buffer and remove detergent as described^18^. Washed beads were resuspended in 50 μl 50 mM NH_4_HCO_3_ containing 20 ng/μl SOLu trypsin (Sigma) and incubated at 25 °C with vigorous shaking overnight. Beads were then pelleted and supernatants were transferred to the new tubes.

### In-solution protein digestion

To identify a protein composition in *E. coli* lysates, proteins from 5 μl of clarified lysates were precipitated with acetone. Four volumes of acetone chilled to −20 °C were added to clarified lysates followed by 1 hour incubation at −20 °C. Precipitates were collected by 10 min centrifugation at 20,000g, 4 °C. Pellets were rinsed with 80% acetone and let dry on air for 15 min. Dried pellets were dissolved in 5 μl 50 mM NH_4_HCO_3_, 8 M urea, then mixed with 50 μl 50 mM NH_4_HCO_3_ containing 20 ng/μl SOLu trypsin (Sigma) and incubated overnight at 25 °C.

After the overnight incubation, digestion reactions were mixed with an equal volume 2% HFBA, incubated at room temperature for 5 min and clarified by 5-min centrifugation at 16,000g. Peptides from supernatants were then desalted using Pierce C18 spin tips (Thermo) according to the manufacturer’s instructions and dried under vacuum. Dried peptides were reconstituted in 0.1% formic acid, concentration was measured at 205 nm on Nanodrop One (Thermo).

### LC–MS analysis of peptides

Depending on concentration, 0.5–2 μg of peptides were analyzed using Orbitrap Fusion Lumos mass spectrometer coupled with Dionex Ultimate 3000 RSLC Nano UHPLC (Thermo). After capturing on 2-cm long (0.2 mm I.D., 5 μm particle size) Acclaim PepMap 100 C18 trap column (Thermo) and washed with buffer A (0.1% formic acid) over 5 min at 5 μl/min, peptides were resolved on a 50-cm long (75 μm ID., 2 μm particle size) EASY-Spray column (Thermo) over 80-min long gradient 2 to 32% buffer B (100% acetonitrile, 0.1% formic acid) at flow rate 0.2 μl/min, followed by steep increase to 90% buffer B over 5 min, and 5-min step elution with 90% buffer B. Data-dependent acquisition method was based on a published protocol ^32^ except that each cycle was set to last for 2 sec instead of 3 sec.

### MS data analysis

Raw MS data were processed using a MAXQUANT 2.0.1 software package ^33^. Protein database included *E. coli* MG1655 proteome (https://www.uniprot.org/proteomes/UP000000625) and a list of common protein contaminants. Search engine was run with default parameters for mass tolerance, up to two missed trypsin cleavages were allowed, variable modifications were methionine oxidation, acetylation of protein N-terminus and cysteine carbamidomethylation. “Second peptide” and “Match between runs” options were enabled. Label-free quantitation was used for MS1-based peptides quantitation.

Further analysis was done in R using quantitation data from a MaxQuant output recorder in “proteinGroups.txt” file. Proteins abundance was normalized to an overall abundance and a protein molecular weight, and log2-transformed. Missing intensities recorded as zero were converted to explicit missing values. Proteins were then ranked by their abundance in a target pull-down, by a difference in the abundance between target and non-specific pull-downs, and by a difference in the abundance between a target pull-down and unfractionated lysate. Next, overall rank ranging from 0 to 1 was calculated as a sum of three ranks. Highly ranking proteins (close to 1) identified in each of three experimental replicates were considered as part of a protein complex involving bait protein—RpoC or CedA.

### Extraction of DNA and preparation of ChIP-seq libraries

Following the immunoprecipitation, washed beads were resuspended in DNA elution buffer containing: 50 mM Tris-HCl pH 8.8, 100 mM NaCl, 10 mM EDTA, 1% SDS and 10 U/ml Proteinase K (NEB), followed by an 18-hour incubation at 56 °C with vigorous shaking. Beads were then pelleted and supernatants were transferred to the new tubes. DNA containing in supernatants was cleaned up using Invitrogen PureLink PCR Purification kit (Thermo). Purified DNA was stored at −20 °C.

NEB Next Ultra II DNA Library Prep kit for Illumina (NEB) was used to prepare DNA libraries according to manufacturer’s instructions. Libraries were amplified using primers supplied with a NEB Next Multiplex Oligos for Illumina kit, Index Primers Set 3 (NEB). Amplified libraries were cleaned up using Sample Purification Beads (NEB) included in the kit. DNA concentration was measured on Qubit (Thermo) and quality analysis was done on TapeStation (Agilent).

### ChIP-seq data analysis

75-bp paired reads were aligned to *E. coli* MG1655 genome (NCBI Reference Sequence: NC_000913.3) using BOWTIE2 (http://bowtie-bio.sourceforge.net/bowtie2/index.shtml). For each genome position, sequencing depth was counted using SAMTOOLS DEPTH tool (https://www.htslib.org). Resulting values were log2-transformed and smoothed using rolling average over a 50-bp wide window. After subtracting background (values obtained from non-specific pulldown) and baseline values (mode of values across all genomic positions), ChIP-seq peaks were called for the positions where local maxima across 50-bp windows had intensity at least 3σ above zero.

### Analysis of differential genes expression using RNA-seq

Bacterial cultures of *E. coli* MG1655 wt, Δ*cedA*::kan and Δ*cedA*::kan bearing paTc-*cedA* plasmid were grown overnight in LB media at 37 °C, 250 rpm. Media was supplemented with 50 μg/ml kanamycin for Δ*cedA*::kan strain, 50 μg/ml kanamycin and 100 μg/ml carbenicillin for Δ*cedA*::kan/paTc-*cedA* strain. Overnight cultures were inoculated at 1/100 vol. into fresh LB containing 10 ng/ml anhydrotetracycline and grown for 2 hours at 37 °C, 250 rpm until the early log phase. From each culture, two 0.5-ml aliquots were taken: control aliquots were mixed with 5 μl H_2_O mQ, and treated with 5 μl 1 mM rifampicin (diluted with H_2_O mQ from 10 mM stock solution in DMSO). After additional 1-hour incubation at 37 °C, 50 μl 10% phenol in ethanol was added to prevent RNA degradation, and bacteria were pelleted by centrifugation at 12000g, 4 °C for 1 min. Media was aspirated and pellets were stored at −20 °C.

Frozen pellets were thawed on ice and resuspended in 50 μl of lysis solution: 50 mM Tris-HCl pH 8.0, 1 mM EDTA, 0.1% Triton X-100, 30 U/μl rLysozyme (Roche), followed by 5 min incubation on ice. Samples were then mixed with 350 μl 1× RNA protection reagent and RNA was purified using a Monarch total RNA extraction kit (NEB) according to manufacturer’s instructions. Purified RNA was stored at −80 °C.

RNA-seq libraries were prepared with NEBNext Ultra II Directional RNA-seq kit for Illumina (NEB) after rRNA depletion (NEBNext rRNA depletion kit by NEB) using 1 μg of total RNA. RNA-seq libraries were amplified and analyzed as described for ChIP-seq libraries.

### RNA-seq data analysis

75-bp paired reads were aligned to *E. coli* MG1655 genome using BOWTIE2. Reads mapping to annotated regions were counted in strand-specific manner using a featureCounts function of Rsubread R package ^34^, including only those reads with the mapping quality above 10 and the length overlap above 0.5. Read counts were then normalized to gene length, transformed to log2 scale and variance stabilizing normalization was applied ^35^. To identify differentially expressed genes, factorial ANOVA test was performed followed by multiple testing correction ^36,37^. Genes were called differentially expressed if associated q-value was below 0.01.

### Next-generation sequencing

The libraries were sequenced on an Illumina NextSeq 500 instrument in 2×75 bp paired end mode. For ChIP-Seq experiments the sequencing depth was typically 10-20 M reads per sample. RNA-Seq libraries were sequenced the depth of 25-30 M reads per sample.

### Proteins expression and purification

Wild-type E. coli RNAP and σ^70^ used in the work were purified as previously descried ^38^. Wild-type E. coli DksA was purified according to ^39^. The open reading frame of the *E. coli* cedA was cloned into the pSUMO vector, which has 10 tandem Histidine tag (10×His) followed by SUMO tag. The plasmid was transformed into *E. coli* strain BL21(DE3) for overexpression and recombinant protein expression was induced with 0.5 mM IPTG when the OD_600_ reached 0.6. After 3 hours at 37 °C, cells were harvested for protein purification. Cell pellets were resuspended in lysis buffer (50 mM HEPES, pH 7.0, 500 mM NaCl, 5% (v/v) glycerol, 15 mM imidazole, 5 mM β-mercaptothanol) supplemented with complete, EDTA-free protease inhibitor cocktail tablets (Roche Applied Science) and lysed using sonication on ice. The cell debris was removed by centrifugation (40,000g for 45 minutes at 4 °C). The supernatant was applied to a HisTrap column (Cytiva) equilibrated in lysis buffer. The column was washed using the lysis buffer until the UV_280_ absorption reaches the baseline.

Protein was eluted with HisTrap Buffer (50 mM HEPES, pH 7.0, 5% (v/v) glycerol, 5 mM β-mercaptoethanol, 250 mM NaCl, 300 mM imidazole). Fractions containing recombinant 10×His-SUMO-CedA were analyzed by SDS-PAGE and then were subjected to tag removal by Ulp1 in dialysis buffer (25 mM HEPES, pH 7.0, 5% (v/v) glycerol, 5 mM β-mercaptoethanol, 100 mM NaCl) at 4 °C overnight. Sample was then applied on a HiTrap SP HP cation exchange chromatography column (5 mL) equilibrated in SP-A Buffer (25 mM HEPES, pH 7.0, 5% (v/v) glycerol, 5 mM β-mercaptoethanol, 100 mM NaCl). The protein was eluted with a linear gradient of NaCl 100 mM to 1 M in 20 column volumes (100 ml). The peak fractions containing CedA were pooled, concentrated, and further purified by a Superdex 75 (10/300) size exclusion column (Cytiva) equilibrated with buffer S (30 mM HEPES, pH 8.0, 200 mM NaCl, 5 mM MgCl_2_, 0.5 mM TCEP). The peak fractions from the Superdex 75 column were pooled, concentrated to 5 mg/ml concentration, flash frozen in liquid nitrogen, and stored at −80°C.

### Nucleic-acid scaffold preparation

Synthetic DNA oligonucleotides containing *ssrA* promoter sequence (−65 to +20) were purchased from Integrated DNA Technologies (IDT). In brief, the nucleic acids were dissolved in RNase-free water (ThermoFisher Scientific) at 1 mM concentration. Template DNA and non-template DNA were mixed at 1:1 ratio, annealed by incubating at 98 °C for 5 min, 75 °C for 2 min, 45 °C for 5 min, and then decreasing the temperature by 2 °C for 2 min until reaching 25 °C. The annealed template DNA: non-template DNA hybrid was stored at −20 °C before use.

### Complex assembly for Cryo-EM

The RNAP holoenzyme (Eσ^70^) was formed by mixing purified RNAP and a 2-fold molar excess of σ^70^ and incubating for 30 min at room temperature. Eσ^70^ was purified on a Superose 6 Increase 10/300 GL column in gel-filtration buffer (30 mM HEPES, pH 8.0, 100 mM KCl, 5 mM MgCl_2_, 0.5 mM TCEP). The eluted Eσ^70^ was concentrated to ≈5.0 mg/mL (≈10 μM) by centrifugal filtration. Annealed *ssrA* promoter DNA was added (3-fold molar excess over RNAP) and the sample was incubated for 30 min at room temperature followed by another 30 min at room temperature after addition of CedA (5-fold molar excess over RNAP). The whole complex was purified on Superose 6 Increase 10/300 GL column equilibrated in gel-filtration buffer. The peak fractions containing RNAP-CedA from the Superose 6 column were analyzed by SDS-PAGE, pooled, concentrated to ≈4 mg/ml for cryoEM sample preparation.

### EM data acquisition

Cryo-grid preparation was performed using FEI Vitrobot mark IV operated at 10 °C and 100% humidity. Aliquots of 4 µl of a freshly purified RPo (≈4 mg/ml), in the presence of 8 mM CHAPSO ^38^, was applied to glow-discharged holey carbon grids (Quantifoil Au, R1.2/1.3, 300mesh). The grids were blotted for 3 sec and flash-plunged into liquid ethane, pre-cooled in liquid nitrogen. The cryo-grids were examined using FEI Arctica operating at 200 KV equipped with K3 direct electron detector, and then good cryo-grids were loaded to FEI Titan Krios electron microscope. All the cryo-images were recorded on Gatan K3 Summit camera operated in super-resolution mode. The magnification is 105,000× corresponding to a final pixel size of 0.852 Å by binning 2 of the original micrographs. For each image stack, a total dose of about 60 electrons were equally fractioned into 42 frames with a total exposure time of 4.2 s. Defocus values ranged from 1.7 to 2.5 μm. In total, 3596 micrographs were collected using Leginon ^40^.

### Image processing

For cryo-EM data sets, beam-induced motion correction was performed using the MotionCorr2 through all frames. The contrast transfer function parameters were estimated by CTFFIND4 ^41^. About 540,000 particles were picked from 3,596 micrographs using Gautomatch in a template-free mode. RELION ^42^ was used for 2D class average and 3D classification. Then 177,841 particles were imported to cryoSPARC ^43^ for final refinement. A reported 2.76 Å map was generated after homogenous refinement and local CTF refinement. All the visualization and evaluation of the map was manipulated within Chimera ^44^, and the local resolution map was calculated using ResMap ^45^.

### Model building and refinement

PDB:6OUL and PDB:2BN8 were used as the template for model building, and *ssrA* promoter sequence was manually placed in COOT ^46^. The resolution was high enough for us to accurately assign the residues of N-terminal of CedA. Phenix ^47^ was used for real space refinement.

### *In vitro* transcription

DNA templates were constructed by PCR amplification and purified from agarose gel using Qiagen gel extraction kit according to the manufacturer.

The sequence of *rrnB*P1, *rrnB*P2 and A1 templates are shown below. The non-transcribed part is italicized. Primers used for PCR amplification are underlined. Bold A and C mark position of the start of transcription for P1 and P2 correspondingly. *rrnB*P1:*<u>tggcagttttaggctgatttggttgaatg</u>ttgcgcggtcagaaaattattttaaatttcctcttgtcaggccggaataactccctat aatgcgcc***A**CCACTGACACGGAACAACGGCAAACACGCCGCCGGGTCAGCGGGGTTCTC CTGAGAA<u>CTCCGGCAGAGAAAGCAAAAATAAATGC</u> *rrnB*P2:*<u>cacgccgccgggtcagcggggttctcctg</u>agaactccggcagagaaagcaaaaataaatgcttgactctgtagcgggaa ggcgtattatgcacac***C**CCGCGCCGCTGAGAAAAAGCGAAGCGGCACTGCTCTTTAACAATT TATCA<u>GACAATCTGTGTGGGCACTCGAAGAT</u> A1:*<u>tccagatcccgaaaatttatcaa</u>aaagagtattgacttaaagtctaacctataggatacttacagcc*ATCGAGAGGGCC ACGGCGAACAGCCAACCCAATCGAACAGGCCTGCTGGTAATCGCA<u>GGCCTTTTTATT</u> <u>T GGATCCCCGGGTA</u>

To measure effect of CedA upon ribosomal promoter initiation, 10 pmol of RNAP were mixed with either CedA up to 5 μM (WT or CedAΔ11N mutant as indicated) or equal amount of TB100A in 60 μL of transcription buffer TB100A (40mM Tris-HCl, pH 8.0, 100 mM NaCl, 10 mM MgCl_2_; 0.1 mg/mL BSA). Samples were incubated 5 minutes at 37 °C and split into two 30-μl parts each. One part from each set was mixed with DksA up to 0.5 μM and ppGpp up to 50 μM, and the second part was mixed with equal amount of TB100A. Samples were incubated at 37 °C for 5 min. Transcription was initiated with addition of 12 pmol *rrnB*P1 PCR promoter fragment pre-mixed with 1 mM ATP and 4 μCi α-[^32^P]-CTP (3,000 Ci mmol^−1^; Perkin Elmer) at 37 °C. 10-μl aliquots were withdrawn at 3, 6, or 12 minute intervals and quenched in fresh tubes with 10 μl Stop Buffer (SB, 1×TBE buffer, 8 M Urea, 20 mM EDTA, 0.025% xylene cyanole, 0.025% bromophenol blue). For *rrnB*P2 promoter, the experiment was performed in the same way except *rrnB*P2 PCR promoter fragment was added together with 100 μM ApC RNA primer, 1 mM GTP and 4 μCi α-[^32^P]-CTP. The products were separated on 23% polyacrylamide gel containing 8 M UREA in TBE (20×20 cm) for 30 minutes at 50 Watt. The gel was transferred onto a film, covered with Saran wrap and exposed to a storage phosphor screen. Screen was scanned on Typhoon Imager (GE) and analyzed using Image Quant software (GE). Relative intensity values were retrieved directly from Image Quant software. Average and standard deviation were calculated in Excel (Microsoft) after normalization and produced less than 10% variation based upon at least three independent experiments.

To measure open promoter complex stability, 10 pmol of RNAP were mixed with equal amount of corresponding promoter DNA in 100 μl of TB100A and then split into two 50-μl aliquots each. The aliquots were mixed with either CedA up to 5 μM or equal amount of TB100A and incubated for 10 minutes at 37 °C. One 10-μl aliquot from each reaction was taken and chased in a new tube with either 1 mM ATP and 1 μCi α-[^32^P]-CTP for *rrnB*P1 or 100 μM ApC RNA primer, 1 mM GTP and 1 μCi α-[^32^P]-CTP for *rrnB*P2 promoter DNA for 5 min at 37°C before quenching with 10 μl SB. The rest of the samples were mixed with heparin (up to 10 μg/ml) and 10 μl aliquots were withdrawn at 1, 2, 4, or 8 min intervals and placed into fresh tube with chase mixtures as described above. Samples were incubated 5 min at 37 °C and quenched with 10 μl SB. Samples were analyzed and visualized as described above.

To measure inhibition with rifampicin in the presence of CedA, 10 pmol of RNAP were mixed with equal amount of A1 PCR fragment in 200 μl of TB100A. Sample was split into two 100-μl parts, one aliquot was mixed with 5 pmol CedA, and the second with equal amount of the same buffer. Samples were incubated for 5 min at 37 °C, and each was split into five 20-μl aliquots. Samples were mixed with rifampicin solution (prepared fresh from dry powder; Sigma) up to 0.2; 0.5, 1 or 2 μg/ml or equal amount of water and incubated for 5 min at 37 °C. RNA synthesis was initiated by addition of 1 mM ATP, GTP, UTP and 10 μM CTP mixed with 0.5 μCi α-[^32^P]-CTP. After incubation at 37 °C, 10-μl aliquots were withdrawn at 3- or 10-min intervals and quenched in new tubes with 10 μl SB. Samples were separated and visualized as described above except 30% polyacrylamide gel was used instead of 23%.

### qPCR

To measure CedA mRNA production, overnight culture of *E. coli* MG1655 was diluted 200 times with 80 mL of fresh LB media and the cells were grown in 500 mL flask at 37°C with shaking until OD_600_ 0.35. Cell culture was split into eight 10-ml parts in 40 ml glass tubes and samples were mixed with corresponding stressors: 2 mM H_2_O_2_, 50 μg/ml Km, 20 μg/ml Gent, 50 μg/ml Ery, 1 μg/ml Cipro and 100 μg/ml Amp. The shaking continued at 37°C for 20 min. Cells were collected by 5-min centrifugation at 5000g, 4 °C. For UV stress measurement, 10 ml of the cell culture was irradiated in a Petri dish with 64 J/cm^2^ UV light (254 nm) at room temperature for 2 min. Cells were allowed to recover in darkness for 10 min at 37°C before collecting.

mRNA was isolated using Master Pure Complete DNA&RNA purification kit (Biosearch technologies) according to the manufacturer’s instructions except that scale was increased 2 times. Resulting RNA was dissolved in TE and concentration was adjusted to 500 ng/ml.

cDNA was produced from 1 μg of total RNA (final cDNA concentration 5 ng/ml) using QuantiTech Reverse Transcription kit (Qiagen) according to the manufacturer’s instructions.

qPCR was performed with 2.5 ng of DNA in triplicates using PowerSYBR Green PCR master mix at Quant Studio 7 qPCR machine (both Applied Biosystems) according to the manufacturer. Primer pairs used for qPCR for *cedA*, *katE* and *gapA* as a reference gene are shown below.

*cedA* or *katE* mRNA levels were normalized to *gapA* mRNA level produced from the same sample and resulting changes in CT values (ΔCT) were compared between treated and untreated samples. −log2 differences in CT values (ΔΔCT) were plotted at Y-axis as fold change.

qRT-PCR primer GapA Forward GCACCACCAACTGCCTGGCT qRT-PCR primer GapA Reverse CGCCGCGCCAGTCTTTGTGA qRT-PCR primer CedA Forward CCGCCAGAACATGCGATAA qRT-PCR primer CedA Reverse GCAGAAATCACTCTCCCATCAG qRT-PCR primer KatE Forward CAGTCACCACTACACGATTCC qRT-PCR primer KatE Revese CTGATTAGTGGTCAGCGCATAA

### Bacterial stress survival assay

Bacterial culture of *E. coli* MG1655 bearing paTc-*cedA* plasmid was grown overnight in LB media supplemented with 100 μg/ml carbenicillin. Next day 10 ml of LB media were inoculated with 20 μL (500× dilution) of the overnight culture and cells were grown at 37 °C with shaking for 1 hour in 40-ml glass tube. Sample was split into two 5-ml parts in 40-ml glass tubes and 100 ng/ml anhydrotetracycline was added to one of them. Cultures were grown for 1 hour at 37 °C with shaking. Samples were again diluted 1000× in 10 ml of fresh LB media containing various stressors: 2 mM H_2_O_2_, 50 μg/ml Kn, 20 μg/ml Gent, 50 μg/ml Ery, 1 μg/ml Cypro. Control cultures with and without induction contained no stressors. Cells were allowed to grow for 90 min at 37 °C with shaking, spun down at 5000g for 5 min, media was discarded and cells were re-suspended in 1 ml of sterile PBS pH 7.2 (Gibco). Cells were serially diluted with PBS and plated at LB-agar plates containing no antibiotics. Plates were grown overnight at 37 °C and colonies were counted.

---

**Extended Data Fig. 1.**
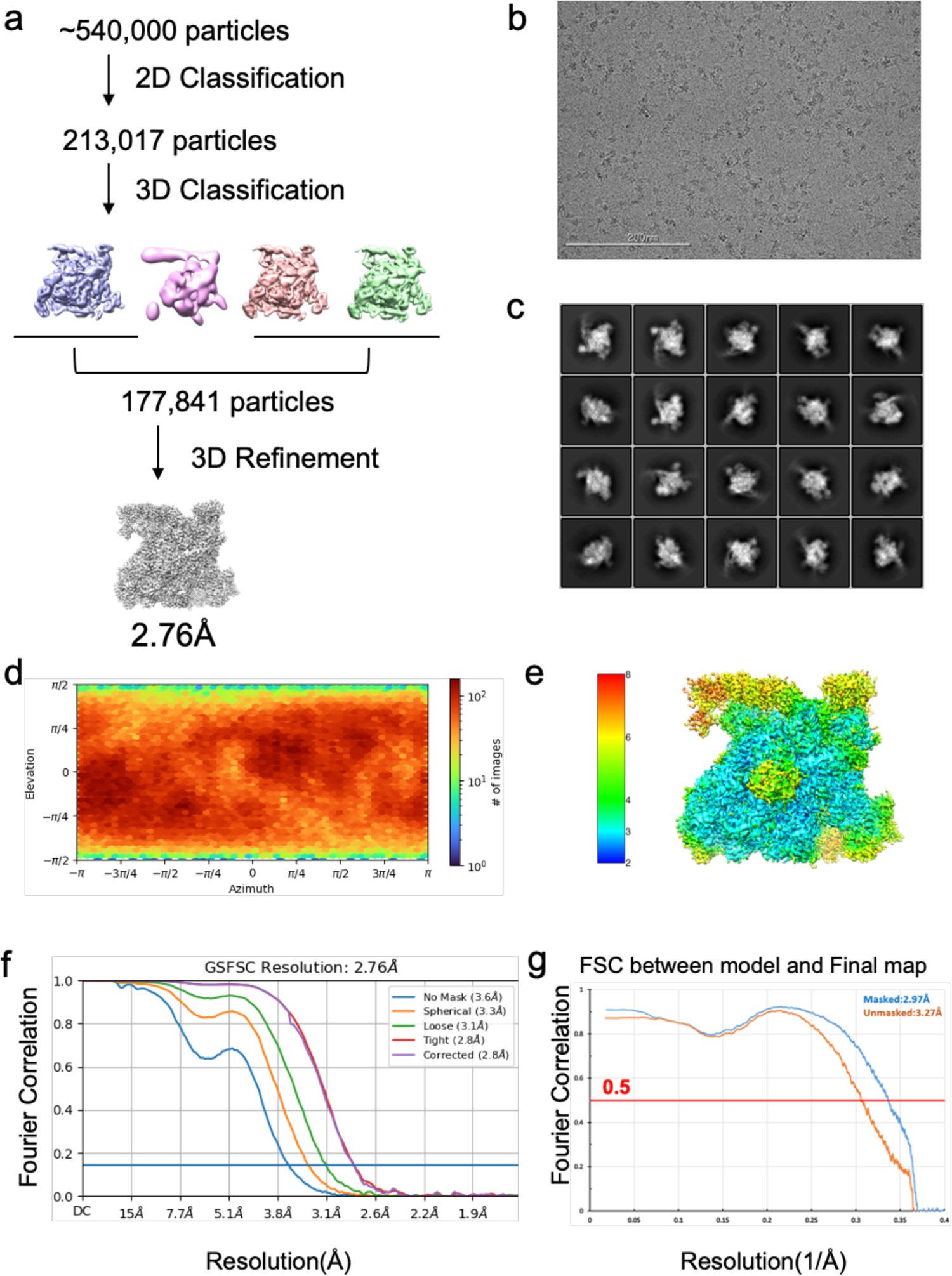
Cryo-EM determination of CedA bound to open-promoter complex. **a**, Flowchart of cryo-EM data processing. **b,** Representative cryo-EM micrographs. Scale bar, 200 nm. **c**, Representative 2D class average shows different views, suggesting randomly distributed particles. **d-f**, Angular distribution, local ResMap estimation and gold-standard Fourier shell Correlation (FSC) curves of final reconstruction. **g**, FSC plots of final reconstruction against the model.

**Extended Data Fig. 2.**
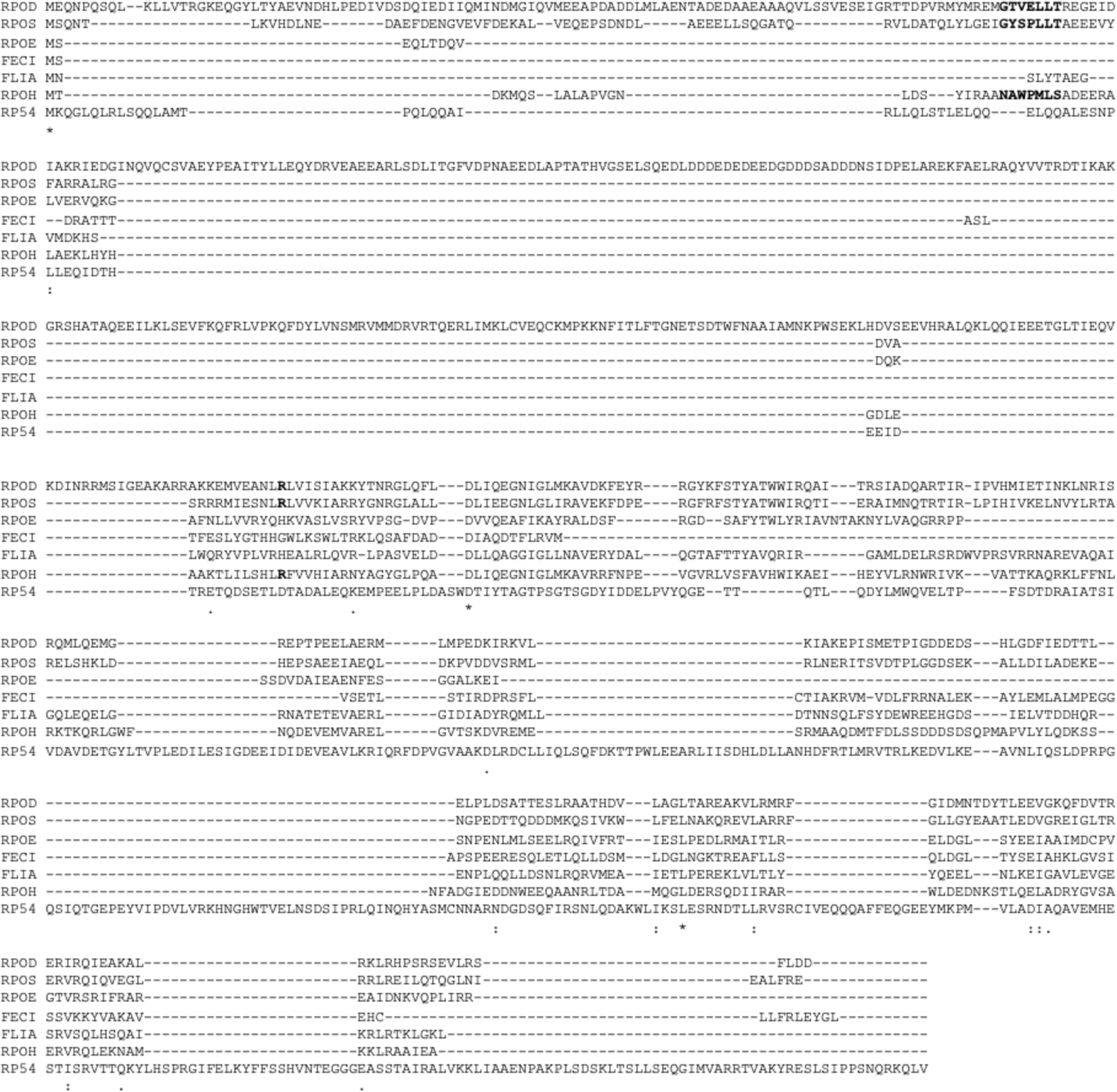
Sequence alignment of *E. coli* sigma factors. Amino acid residues participating in interaction with CedA are shown in bold.

**Extended Data Fig. 3.**
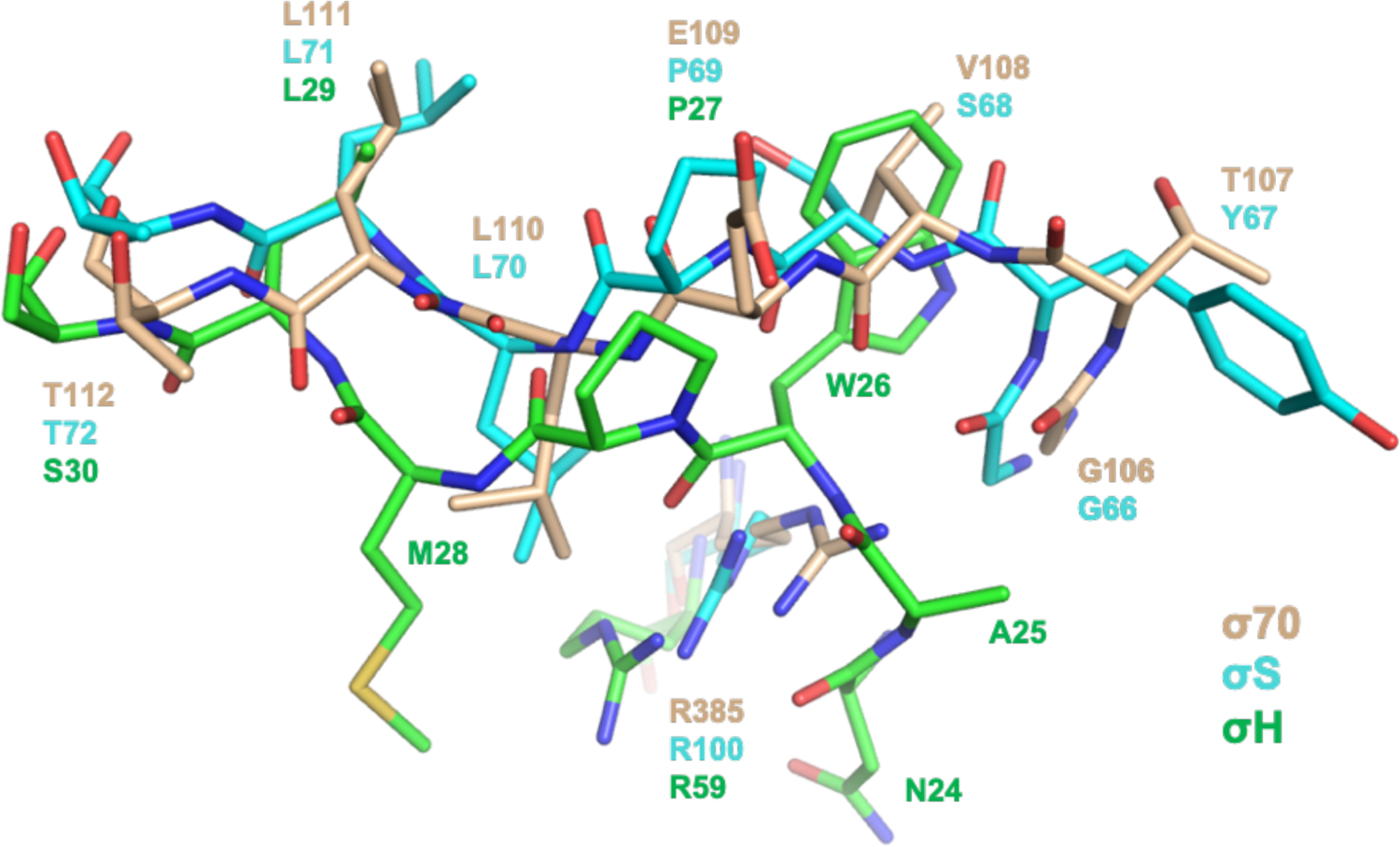
Structural comparison of region 1.2 in σ^70^, σ^S^ and σ^H^. Structure models of σ^70^ (shown in wheat, this study), σ^S^ (shown in cyan, PDB: 6OMF), and σ^H^ (shown in green, AlphaFold: AF-P0AGB3-F1) were aligned using PyMol 2.4. Region 1.2 interacting with CedA is shown.

**Extended Data Fig. 4.**
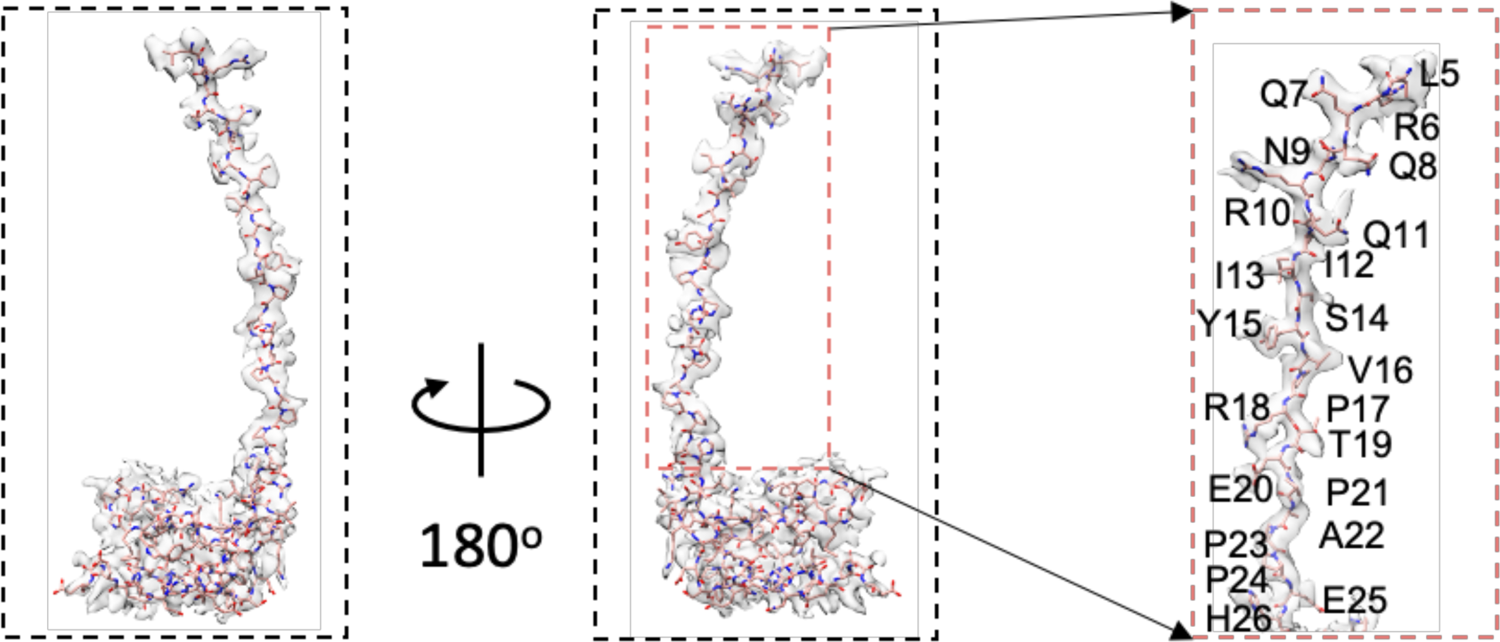
Cryo-EM density of CedA extracted from 2.76 Å cryo-EM map.

**Supplementary Table 1.** Proteins co-isolated with RNA polymerase.

**Supplementary Table 2.** Proteins co-isolated with RNA polymerase, sorted.

**Supplementary Table 3.** Proteins co-isolated with CedA, sorted.

**Supplementary Table 4.** CedA ChIP-seq data.

**Supplementary Table 5.** RNA-seq data for WT, ΔCedA with or without Rif treatment, with or without pCedA

**Supplementary Table 6.** Oligonucleotides used for strains constructed in this work.

